# Deep Learning For Denoising Hi-C Chromosomal Contact Data

**DOI:** 10.1101/692558

**Authors:** Max Highsmith, Oluwatosin Oluwadare, Jianlin Cheng

## Abstract

**Motivation:** The three-dimensional (3D) organization of an organism’s genome and chromosomes plays a significant role in many biological processes. Currently, methods exist for modeling chromosomal 3D structure using contact matrices generated via chromosome conformation capture (3C) techniques such as Hi-C. However, the effectiveness of these methods is inherently bottlenecked by the quality of the Hi-C data, which may be corrupted by experimental noise. Consequently, it is valuable to develop methods for eliminating the impact of noise on the quality of reconstructed structures.

**Results:** We develop unsupervised and semi-supervised deep learning algorithms (i.e. deep convolutional autoencoders) to denoise Hi-C contact matrix data and improve the quality of chromosome structure predictions. When applied to noisy synthetic contact matrices of the yeast genome, our network demonstrates consistent improvement across metrics for contact matrix similarity including: Pearson Correlation, Spearman Correlation and Signal-to-Noise Ratio. Positive improvement across these metrics is seen consistently across a wide space of parameters to both gaussian and poisson noise functions.

## Introduction

The three-dimensional (3D) structure of the genome has been shown to play a significant role in many biological processes such as gene regulation [1], methylation [2], and gene expression [3]. Two of the most prolific strategies for analysis of 3D chromosome structure are fluorescent in situ hybridization (FISH) [4][5] and chromosome conformation capture (3C) techniques [6].

FISH utilized fluorescent probes which bind to parts of the chromosome with high sequence complementarity and can be used to measure distances between genomic regions, However, FISH is heavily limited by its low resolution and low throughput.

Chromosome capture technologies, such as Hi-C [7], use high-throughput sequencing to find the nucleotide sequence of fragments in spatial proximity, which are then used to construct lists of contact fragments. The data obtained from Hi-C techniques can be used to construct contact matrices where entries indicate contact between genomic regions and the number of nucleotides contained in one equal-sized region, called a bin, is referred to as the resolution.

A variety of methods have been developed to use Hi-C contact matrices to construct 3D structures such as [8] LorDG, [9] 3DMax, [10] Shrec3d, [11] Mogen, and [12] Pastis. However the effectiveness of any model is inherently bottlenecked by the quality of the data provided as input. The work from [13] Yaffe et al shows that several forms of systematic bias affect Hi-C experimental procedures such as GC content of trimmed ligation junctions, sequence uniqueness and distance between restriction sites. In addition to recognized forms of bias, there is always potential for unidentified experimental noise and bias inhibiting the quality of data.

The concept of utilizing deep neural networks as a preprocessing tool to improve the quality of Hi-C data was first proposed by Yan et al in 2018 [14]. Their network, HiCPlus, used a convolutional neural networks to infer high resolution Hi-C data from experimentally obtained low resolution Hi-C data. By training on low resolution Hi-C datasets with higher resolution Hi-C matrices as labels they obtained comparable maps using 1/16th of the needed experimental reads. Since then the network HiCNN has improved upon Hi-C plus by utilizing a deeper architecture[15].

The deep neural networks discussed in this work differ from both HiCPlus and HiCNN in two key ways: First the enclosed experiments do not seek to raise the resolution of experimentally obtained Hi-C data, but rather aims to remove experimental noise while maintaining the same contact map resolution. Second, the network considered is a denoising autoencoder, a flavor of unsupervised learning, rather than the supervised deep network used in HiCNN and HiCPlus. Therefore, our method does not require labeled data, which is very rare in this domain, to train the deep network. Thus, our method can be applied to denoise any Hi-C data. Our experiment demonstrates the method can consistently improve the quality of chromosomal contact matrices derived from Hi-C data.

## Method

Autoencoders are a class of neural network which utilise an unsupervised learning algorithm to encode data from a sample space into a (usually lower dimensional) latent space in such a manner that reconstructions which closely resemble the original data sample can be generated from the encoding. Generally speaking an autoencoder with sample space *X* and latent space *Z* applied to a dataset *D* = {*x*(1), .., *x*(*k*), ..*x*(*K*)} lis composed of 3 parts.

1. An encoding function *φX* – > *Z* which maps data samples to latent space, *φ*(*x*(*k*)) = *z*(*k*)
2. A decoding function *ψZ* – > *X* which maps latent space representations back to the sample space *ψ*(*k*)) = *x*_*r*_(*k*)
3. A learning algorithm *A*(*φ,ψ,D*) which minimizes some loss function *L*(*x*, *x*_*r*_)

Autoencoders have been shown to be useful across many domains including: feature learning, clustering and denoising.

In traditional autoencoders, the loss function *L* is minimized with the target reconstruction being the same data vector as the sample input. With Denoising Autoencoders the initial input is perturbed via a noise function prior to passing through the autoencoder [16]. The learning algorithm is then applied using the clean data as the target for reconstruction thus the loss function,*L*(*x*_*noise*_, *x*_*clean*_, is minimized. The hypothesis being that by training in this mechanism the information loss during the dimensionality reduction will be the noise rather than the true signal.

Our experiments begin with a collection of clean and noisy matrices in Hi-C contact space, denoted *x*_*i,clean*_ and *x*_*i,noisy*_, related by the noise inducing function *η* such that *x*_*i,noisy*_ = *η*(*x*_*i,clean*_). In the ideal we aspire for a function *η*^−1^ which perfectly removes all noise mapping *x*_*i,noisy*_ back to *x*_*i,clean*_. However, because the noise function *η* is not constrained to be injective or surjective, the existence of a well defined inverse is not necessarily guaranteed. Consequently in this paper we limit ourselves to the task of developing a function *A*(*x*_*i,clean*_) which approximates *η*^−1^, and whose characterization is determined by a neural network.

When denoising Autoencoders are applied to the task of image denoising, the learning algorithm is traditionally trained using the clean data as labels, i.e *L*(*x*_*noise*_, *x*_*clean*_) [16]. However, in the task of denoising Hi-C contact matrices, true clean labels are not always available, hence the need for denoising. To capture this scenario we also train the autoencoders in a purely unsupervised fashion, i.e. *L*(*x*_*noise*_, *x*_*noise*_): in hopes that the information lost post training will primarily be noise rather than signal. For the remainder of the paper we will refer to these two variants of the denoising autoencoder training strategy as semi-supervised and fully-unsupervised respectively.

**Figure 1.**
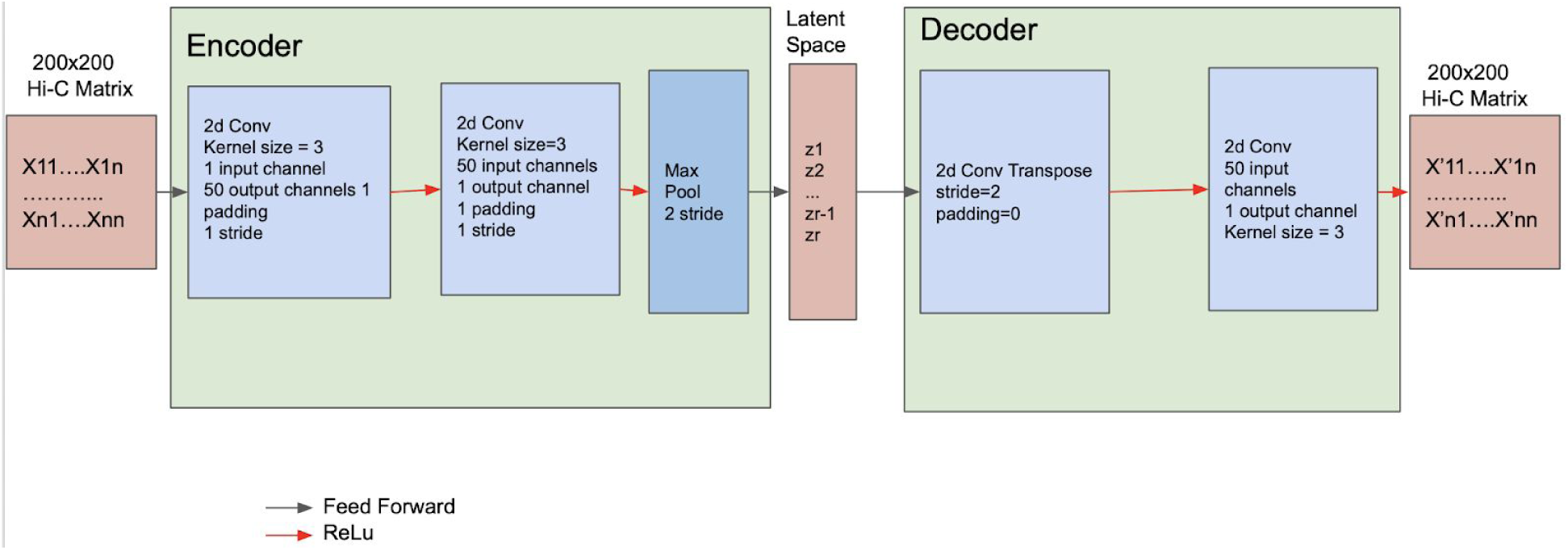
The architecture of the deep autoencoder for denoising Hi-C chromosomal contact matrices. The encoder consists of two convolutional layers and one max pooling layer. The decoder has two convolutional layers. ReLu function is applied to the convolutional layers.

### Deep Convolutional Autoencoder

Our network is a feedforward convolutional neural network with symmetric encoders and decoder. We select latent spaces to be half the size of the input space and give both encoders and decoders two convolutional layers. We downsample to the latent space via max pooling and upsample via a trainable convolutional transpose. Each layer is separated by a ReLU activation function. We implement all networks using Pytorch [18]. We train all networks with the adam optimizer. We also append the network with a format filter which ensures that the diagonals of our interaction matrices remain 0, and enforce symmetry by casting the lower triangular matrix to be a symmetric reflection of the upper triangle matrix.

We train using a custom loss function differing in the purely unsupervised and semi-supervised instances.

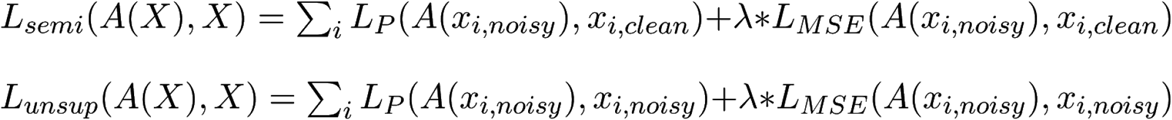

Where

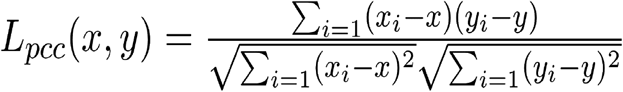

Maximizes the pearson correlation between the output and target matrices and

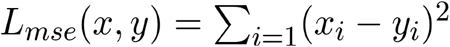

Is the standard mean squared error loss. The variable *λ* is used as a term-weighting hyperparameter. Both of these components of the loss function are applied because correlation and squared error are used in evaluating similarity of three dimensional structure in many 3D conformation predicting tools [8][9]. These metrics, along with others used in evaluation, are further explained in the Evaluation Metrics section.

## Results

To evaluate the potential effectiveness of our Denoising Network we conduct experiments on both synthetic and publicly available experimentally obtained Hi-C datasets.

### Synthetic Datasets

We simulate synthetic chromosomal contact datasets from the theoretical 3D model of the yeast genome [19]. The supplementary material from Duan et al provides a 3D structure of the yeast genome at kilobase resolution. Keeping the full resolution of the original yeast structure, the genome is split into a collection of smaller, non-overlapping fragments with equal bead counts. The Hi-C experiment is then simulated to form 5 contact maps from each fragmented structure, while introducing the noise function *η* Figure 2. This is done for bead counts of 150 and 75 forming datasets containing 850 and 1730 samples respectively. We then seperate each dataset by 80% 10% 10% training, validation, testing split as shown in Figure 2.

**Figure 2:**
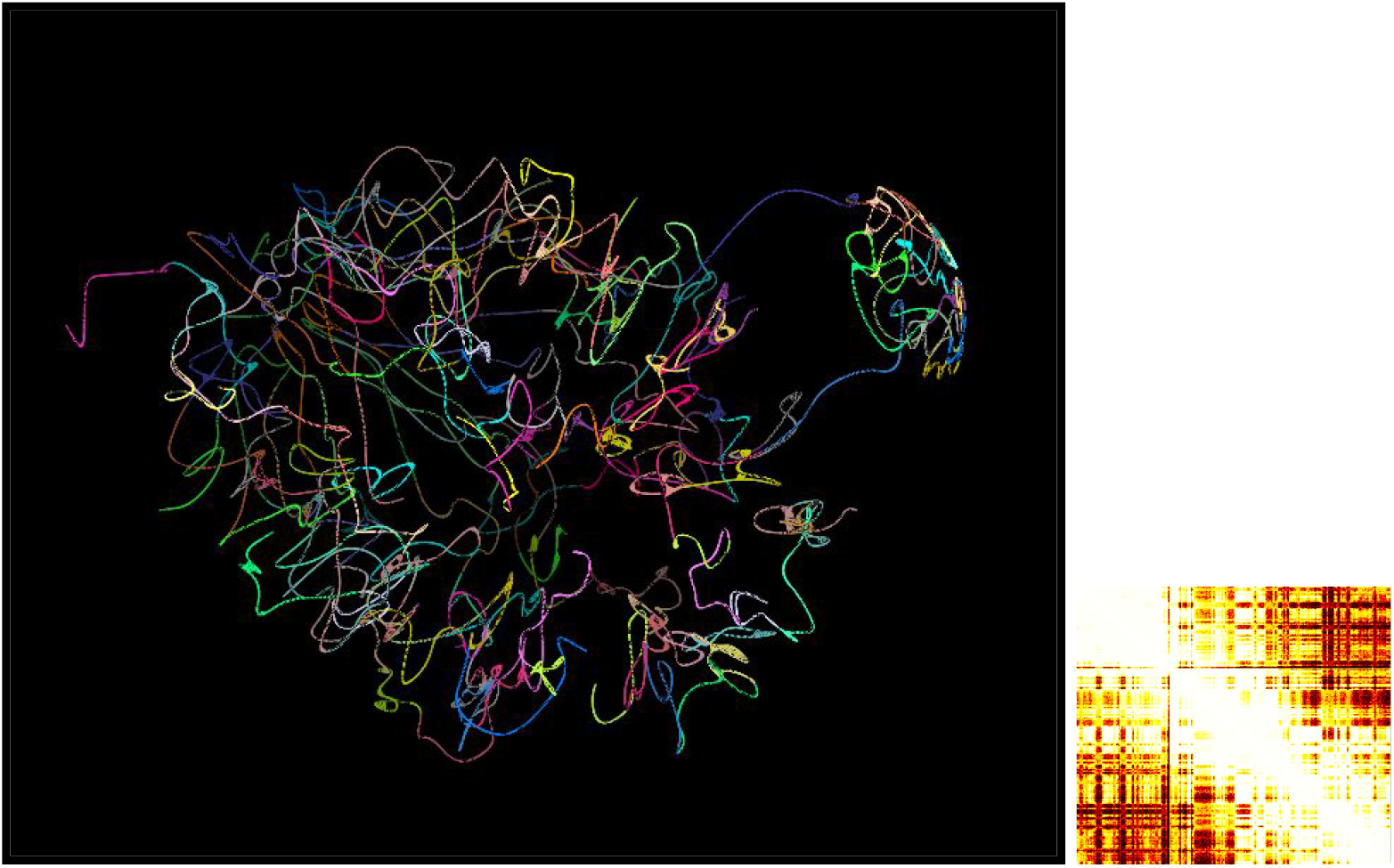
(left) 3d structure of the yeast genome. Each color change corresponds to a fragment used to generate a 150×150 Hi-C contact matrix (right).

For the synthetic datasets, noise is introduced into these contact matrices as a function of distance by the following equation

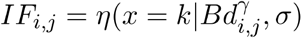

Where *IF*_*i,j*_ is the interaction frequency (i.e. contact count) between beads *i* and *j* in contact matrix space, *d*_*i,j*_ is the distance between bead *i* and *j* in the ground truth conformation. The variable *γ* is set to −1 as shown in Trieu et al [8], rather than −3 as is shown in Varoquaux et al [12] to remain consistent with the fractal rather than equilibrium model of the genome [20]. The variables B and *σ* are used as parameters to influence the level of noise introduced by *η* and the selection of probability function *η* determines the nature of noise considered in an individual experiment. Prior to the introduction of noise the value *d*_*i,j*_ is divided by the average of all distances so as to mitigate errors due to rounding.

We consider two parametric families of univariate distributions for the noise function *η*: Poisson and Gaussian.

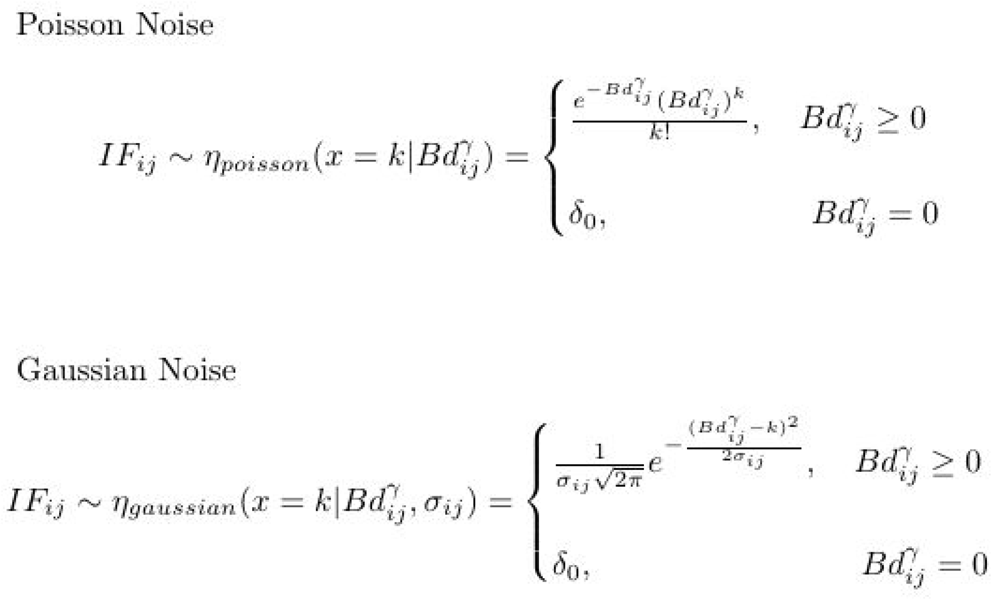

Both noise functions represent potential variance in contact frequency between two genomic regions, while preserving the assumption that as 3D spatial proximity increases the contact count will also increase in a relatively monotonic nature. The poisson noise function is the noise function used in [8] and [12], where an increase in parameter B causes the Signal Noise Ratio to increase due to decreased incidence of lost information in rounding errors. One fundamental characteristic of a poisson function is that its mean and variance are necessarily the same value. Thus, in the context of noise introduction if *d*_*i,j*_ is increased the variance in values for *IF*_*i,j*_ must also increase in a strictly linear fashion. It is unclear if this dependency is inherent to the biological process being modeled. Use of the gaussian noise function allows us to remove this inherent dependency and instead constrain the variance to be constant across all components of the contact matrix.

In image denoising literature levels of noise are often quantified by the Signal Noise Ratio [21]. As its name implies the SNR of a noisy image is the ratio between the magnitude of a signal and the difference between the true signal and its noisy representation. An interpretation of SNR in the context of Hi-C Contact Matrix determination is provided by Trie et al [11] via the equation

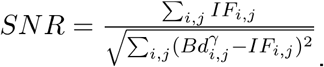

A high SNR is indicative of a low level of corruption by noise. By determining the SNR of the noisy matrices created by each respective noise function we can contrast the effects of the different parametric families of noise functions on model accuracy.

### Real Hi-C Datasets

We run experiments using real Hi-C data obtained through procedures on the GM12878 cell line and available at the GEO GSE63525 [22]. All datasets used were downloaded in .hic format and passed to the GenomeFlow tool [9] to generate contact matrices readable by our network. Rather than inserting synthetically manufactured noise into the contact matrices, we consider matrices generated by two different procedures, one of which has been demonstrated to provide higher quality data than the other. We treat the lower quality data as the noisy set and the higher quality data as the clean set

To construct the noisy contact matrices we used the hi-c data generated via the Dilution protocol originally proposed by Liebermann et al [7]. To construct the clean contact matrices we used the more modern In Situ protocol described by Rao et al [22]. Defining map resolution as the lowest number of bases per bin satisfying the condition that 80% loci have greater than 1k contacts, the two data sets have map resolution 1kb and 1Mb respectively. This illustrates that although the In Situ contact matrices are not ‘ground truth’ they are considerably higher quality than the Dilution contact matrices, and can be used as benchmarks for denoising of the lower quality Dilution contact matrices.

We run experiments on contact matrices binned at the matrix resolution of 25kb. We chose this resolution because at lower resolutions (50kb, 100kb) the correlation between the noisy and clean matrices was already very high. We constrain our experiments to contact matrix fragments of size 200bin x 200bin centered along the chromosome diagonal. For the semi-supervised experiments we use chromosomes 12, 11, and 17 as training, validation and testing sets yielding sets containing 2477, 2499, and 1424 samples respectively. On the fully-unsupervised experiments we use samples from chromosome 12.

### Evaluation Metrics

Remaining consistent with the existing literature[8][9][10][11][12] We use two key metrics for evaluation of contact matrix similarity: Pearson Correlation Coefficient and Spearman Correlation Coefficient.

Pearson Correlation Coefficient:

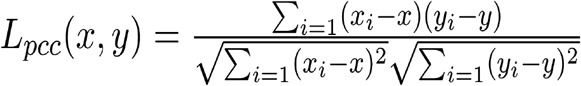

Spearman Correlation Coefficient:

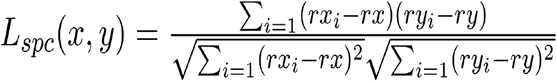

Where rx and ry are the rank variables.

We refer to the Pre and Pas values of a metric as the comparison of noisy data with clean data and the comparison of denoised data with clean data respectively. For an effective denoising network our Pre values will be smaller than Pas values for correlation metrics and Pre values will be greater than Pas values for the Error metric. The training process of our network deals with matrices in contact space so we evaluate the effectiveness of our model using noisy, denoised and clean contact matrices.

### Performance on varying noise parameters with synthetic data

We see from Figure 2 that the Deep Convolutional Autoencoder shows fairly consistent improvement on SNR, SPC and PCC respectively across the grid of Gaussian noise parameters. As would be expected increases of the variance parameter *σ* and decreases of the sampling parameter *β* show degradation of correlation in the noisy contact maps. The quality of denoised data continues to improve across all entries of the parameter grid for spearman correlation, and the level of improvement is proportional to the quality of the noisy data. For the PCC metric improvement is fairly consistent however at the edges of the parameter grid there are instances where there is degradation as a consequence of denoising. However, it is worth noting that this only occurs in instances where the PCC metric is already greater than .95, hence the noisy data has a very high SNR already, leaving little room for improvement.

**Figure 2:**
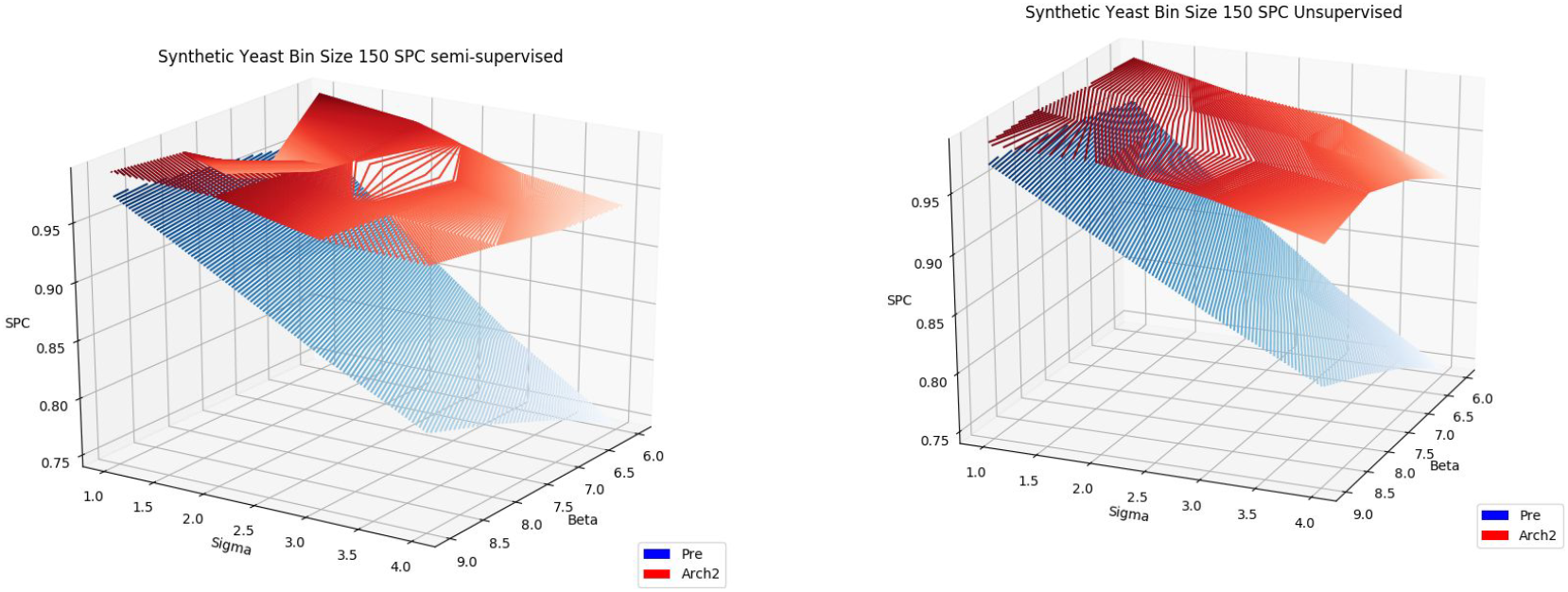
Spearman’s Correlation (SPC) for Gaussian noise on semi-supervised (left) and fully-unsupervised (right) experiments Red surface displays correlation with clean matrix after denoising, Blue indicates correlation correlation prior to denoising

**Figure 3.**
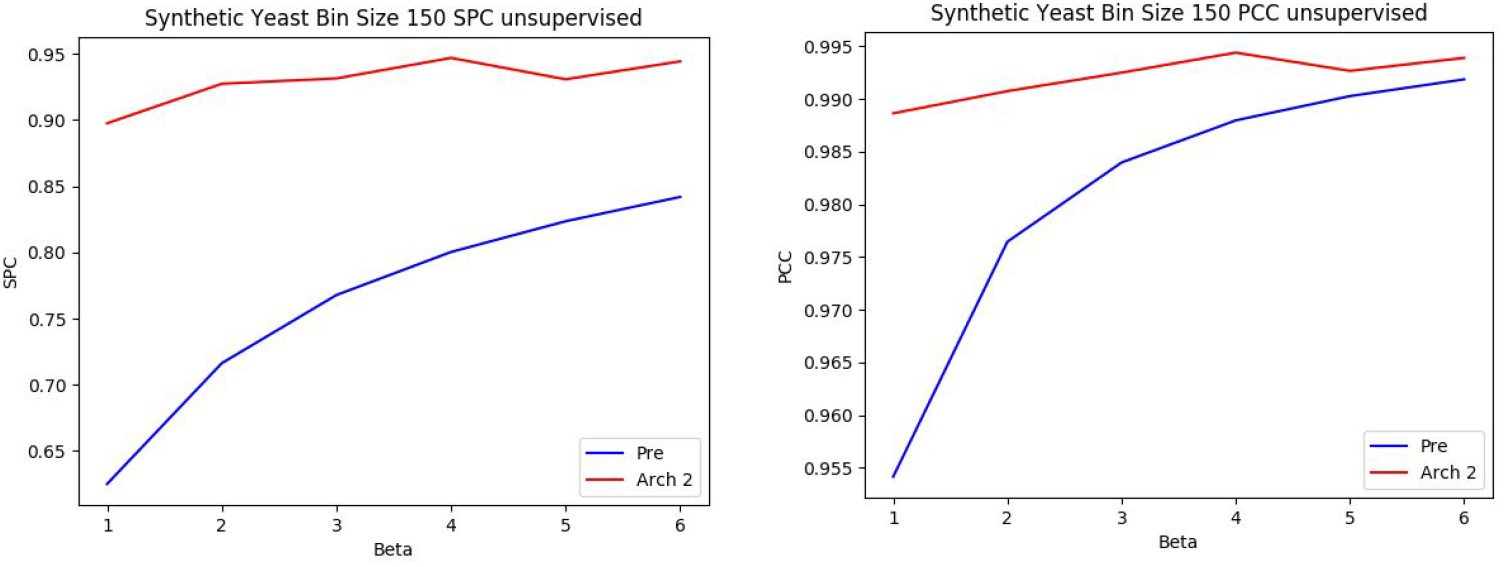
The SPC (left) and PCC (right) metrics for denoised data in unsupervised experiments with Poisson noise

The Deep Convolutional Autoencoder shows consistent improvement for SNR, SPC and PCC metrics across the entire tested grid of Poisson parameters. For unsupervised experiments both the SPC and PCC improvement remains positive but decreases proportionally as noise decreases, likely because there is less room for improvement. As with the Gaussian experiments we see the PCC is very resilient to noise while SPC is more significantly affected. The network show enormous gains in SPC for very noisy data at *beta* = 1 rising from 0.68 SPC to 0.91 SPC

### Performance on Real Hi-C Chromosomal Contact Data

We see from Table 1 that the convolutional autoencoder displays consistent improvement along the SPC metric in both the semi-supervised and fully-unsupervised experiments. PCC also shows substantial improvement in the semi-supervised experiments while showing negligible change in the fully-unsupervised experiments. Consideration of the minor decrease in performance on the PCC metric for the fully-unsupervised experiment should take the following into account. First, the PCC is already fairly high between the Dilution and In Situ training sets, and even after the minor drop, the two matrices are still highly correlated along this metric. Secondly, PCC imposes a linearity constraint which the SPC metric circumvents and which may not be grounded in biological truth. Third the SPC metric is more prolific throughout the literature. In the semi-supervised experiments the PCC metrics shows significant improvement increasing correlation by a full 0.07 in test correlation. The consistent increase in performance along the SPC metric in both semi-supervised and fully-unsupervised experiments demonstrate network effectiveness at denoising real Hi-C data.

**Table 1.** SPC and PCC of Unsupervised and Supervised experiments on GM12878 Cell Line. X’s indicate no measurement. Unsupervised experiment have no labels and thus the training set is the testing set. Validation for model selection is always done using SPC metric as it is more prolific in Hi-C literature

| Experiment | Training Pre | Training Pass | Val Pre | Val Pass | Test Pre | Test Pass |
| --- | --- | --- | --- | --- | --- | --- |
| Semi-Supervised SPC | 0.9066 | 0.9274 | 0.9 | 0.919 | 0.8838 | 0.895 |
| Unsupervised SPC | 0.9066 | 0.9148 | x | x | x | x |
| Semi-Supervised PCC | 0.9199 | 0.9788 | x | x | 0.8077 | 0.8709 |
| Unsupervised<br>PCC | 0.9199 | 0.9116 | x | x | x | x |

### Comparison to Non-Deep learning based methods

We benchmark our denoising of the synthetic yeast dataset against two prolific methods in image denoising literature, the gaussian filter and wavelet denoising. The implementations of both of these methods come from packages in scikit learn [24]. These benchmark methods are first applied to the noisy contact matrices. Because these methods are used in generic image denoising they do not naturally impose format constraints for hi-c matrices. Thus prior to comparison with the Deep Autoencoders, wave and gauss matrix diagonals are set to 0 and symmetry is imposed by reflection of the upper triangular matrix across the diagonal. Table 2 shows that both the unsupervised and semi-supervised deep convolutional autoencoders perform better at the task of denoising than either of these methods. In this case the wavelet denoising strategy actually results in a decrease in performance across both metrics. The poor performance of these denoising benchmarks may be due to the gaussian filter and wavelet denoising typical intended application being in the realm of making images appear less noisy rather that the specific application of preserving chromosome contact information.

**Table 2.** Comparison of Deep Denoising Network to non-deep learning based methods (Gaussian Filter and Wavelet Denoising).

| Yeast<br>Gaussian<br>Sigma 1 Beta 5<br>Test Data | Pre | Gaussian Filter | Wavelet<br>Denoising | Pas Deep<br>Conv<br>Unsupervised | Pas DeepConv<br>Semi-Supervised |
| --- | --- | --- | --- | --- | --- |
| Matrix SPC | 0.8003 | 0.8961 | 0.79905 | 0.933 | 0.934 |
| Matrix PCC | 0.9879 | 0.8449 | 0.8919 | 0.993 | 0.993 |

## Conclusion

Hi-C data can be used to build contact matrices which in turn are used by different tools to study enhancer-promoter interactions and describe the 3D conformation of Chromosomes and Genomes. In this paper we use a convolutional denoising autoencoder for the task of denoising Hi-c data. We demonstrate our proposed networks effectiveness at denoising data in both a fully-unsupervised and semi-supervised setting on both syntheticlly constructed and real Hi-C datasets.

